# Bioluminescent zebrafish transplantation model for drug discovery

**DOI:** 10.1101/2022.03.03.482856

**Authors:** Martina Hason, Jovana Jovicic, Ivana Vonkova, Milan Bojic, Theresa Simon-Vermot, Richard M. White, Petr Bartunek

## Abstract

In the last decade, zebrafish have accompanied the mouse as a robust animal model for cancer research. The possibility of screening small-molecule inhibitors in a large number of zebrafish embryos makes this model particularly valuable. However, the dynamic visualization of fluorescently labeled tumor cells needs to be complemented by a more sensitive, easy, and rapid mode for evaluating tumor growth *in vivo* to enable high-throughput screening of clinically relevant drugs. In this study we proposed and validated a pre-clinical screening model for drug discovery by utilizing bioluminescence as our readout for the determination of transplanted cancer cell growth and inhibition in zebrafish embryos. For this purpose, we used NanoLuc luciferase, which ensured rapid cancer cell growth quantification *in vivo* with high sensitivity and low background when compared to conventional fluorescence measurements. This allowed us large-scale evaluation of *in vivo* drug responses of 180 kinase inhibitors in zebrafish. Our bioluminescent screening platform could facilitate identification of new small-molecules for targeted cancer therapy as well as for drug repurposing.

## INTRODUCTION

Over the last five decades, there has been a significant amount of resources and efforts invested into cancer research (Hanahan and Weinberg, 2000; Hanahan and Weinberg, 2011). Along with genetically engineered models, tumor cell transplantation is an additional suitable method for the assessment of cancer cell engraftment, invasiveness and treatment possibilities *in vivo* (Baeten et al., 2019; Capasso et al., 2019; McCune et al., 1988; Patton et al., 2021b). While the mouse is a powerful and widely used animal model, it is relatively difficult to accomplish high-throughput small molecule inhibitor screening in this model. For more than a decade, zebrafish (*Danio rerio*) has accompanied the mouse as another promising model animal for human tumor xenograft assays (Cagan et al., 2019; Kirchberger et al., 2017; Topczewska et al., 2006). Drug discovery connected to inhibitor screening is a field where zebrafish have a great potential (Bowman and Zon, 2010; MacRae and Peterson, 2015; Patton et al., 2021b). With the small-sized transparent embryos lacking the adaptive immune system, it is feasible to graft and track high numbers of experimental animals in a relatively short time. Further, its high evolutionary conservation of genes connected to cancerogenesis makes zebrafish a flexible animal for human disease modeling (Fazio et al., 2020; Howe et al., 2013; Lam et al., 2006).

Zebrafish embryos are naturally transparent and can engraft transplanted cancer cells until around the seventh-day post fertilization when the adaptive immune system starts to mature. This allows for simple *in vivo* tracking and imaging of fluorescently labeled cancer cells and the study of early tumor growth and dissemination with the involvement of the tumor microenvironment (Hason and Bartunek, 2019; Povoa et al., 2021; Tang et al., 2016; White et al., 2013). For longer-term studies, it is possible to use genetically immunocompromised animals, such as the *rag2*^E450fs^ or the *prkdc*^D3612fs^ mutant lines (Tang et al., 2014; Tang et al., 2016; Yan et al., 2019; Yan et al., 2020).

Small-molecule pharmacological screening in zebrafish embryos can be carried out with medium to high throughput (Colanesi et al., 2012; MacRae and Peterson, 2015; Patton et al., 2021b; Richter et al., 2017). Targeting of cancer-related pathways in zebrafish embryos assisted in the discovery of compounds such as leflunomide (White et al., 2011), Lenaldekar (Ridges et al., 2012), perphenazine (Gutierrez et al., 2014), and clotrimazole (Precazzini et al., 2020) as potential treatment options for cancer mono- and combined therapy of human leukemia and melanoma. Even though there are many established zebrafish xenograft models, up until now, there were only a few larger inhibitor screens done in zebrafish allograft or xenograft models (Almstedt et al., 2021; Fazio et al., 2020; Fior et al., 2017; Somasagara et al., 2021; Tulotta et al., 2016; Wertman et al., 2016). This is likely due to the limited options of workflow automation in transplantation studies. In the field of zebrafish transplantation studies, where mainly fluorescent cell lines are used for imaging purposes, we decided to take steps towards the utilization of a different readout mode.

NanoLuc^®^ luciferase (NanoLuc), a small luciferase subunit derived from a deep-sea shrimp *Oplophorus gracilirostris* (Hall et al., 2012; Schaub et al., 2015), enabled us to easily track the number of cancer cells *in vivo*. Bioluminescence has been reliably used in mice as well as zebrafish for tracking cell fates *in vivo* (Astuti et al., 2017; de Latouliere et al., 2021; Manni et al., 2019; Stacer et al., 2013). Because of the ease of measuring luminescence in real-time *in vivo*, this setup is suitable for high-throughput screening. Bioluminescence-based analysis ensures higher sensitivity, less background and therefore accurate cancer cell growth quantification compared to the conventional fluorescence-based readout. Here, we established a bioluminescent small-molecule screening system which allowed us to evaluate kinase inhibitors in zebrafish transplantation models of melanoma and myeloid leukemia. We found inhibitors targeting cell proliferation, migration and survival as hits in our *in vivo* screen. With our work, we show that zebrafish can serve as a robust pre-clinical screening model for the discovery of new cancer therapeutics.

## RESULTS

### Establishing a bioluminescent platform for tumor cell transplantation in zebrafish

To develop a zebrafish transplantation model with bioluminescence as a readout we prepared two cancer cell lines with the concurrent expression of fluorescent proteins and NanoLuc. NanoLuc is the brightest and the most stable luciferase available, therefore we chose it for our experiments (Hall et al., 2012). We labeled the zebrafish melanoma cancer cell line ZMEL1 with a double reporter system containing *EGFP* and *NanoLuc* (Fig. S1A). These cells were sorted by fluorescence-activated cell sorting (FACS) to obtain single clones according to the expression level of *EGFP*. Further, we prepared cells of the human chronic myelogenous leukemia (CML) cell line K562 in a similar way, expressing *mCherry* and *NanoLuc* (Fig. S1B). We confirmed the insertion of the reporter into the genome of cells by sequencing and tested the expression and activity of luciferase by *in vitro* luciferase assay. We were able to distinguish luminescence with high sensitivity, to the level of single cells (Fig. S1A-B). After confirming that our double-reporter worked *in vitro*, we moved to cell transplantation into zebrafish embryos. We found the Duct of Cuvier as the best site for transplantation into the bloodstream of 2 days post fertilization (dpf) *casper*, *prkdc*^−/−^ embryos (Moore et al., 2016). Transplanted embryos were imaged at 3 dpf and distributed into 96-well plates for *in vivo* luminescence assay. We observed no signs of toxicity associated with Furimazine, the substrate of NanoLuc. We plotted the measured luminescence in single embryos against the relative amount of fluorescence to determine correlation of the two readouts (Fig. 1A). Indeed, for both cell lines there was a positive correlation of *EFGP* and *NanoLuc* in ZMEL1 as well as of *mCherry* and *NanoLuc* in K562 (Fig. 1A). Therefore, we used luminescence as our readout for measuring cancer cell growth and its inhibition *in vivo* in all our further experiments.

**Fig. 1.**
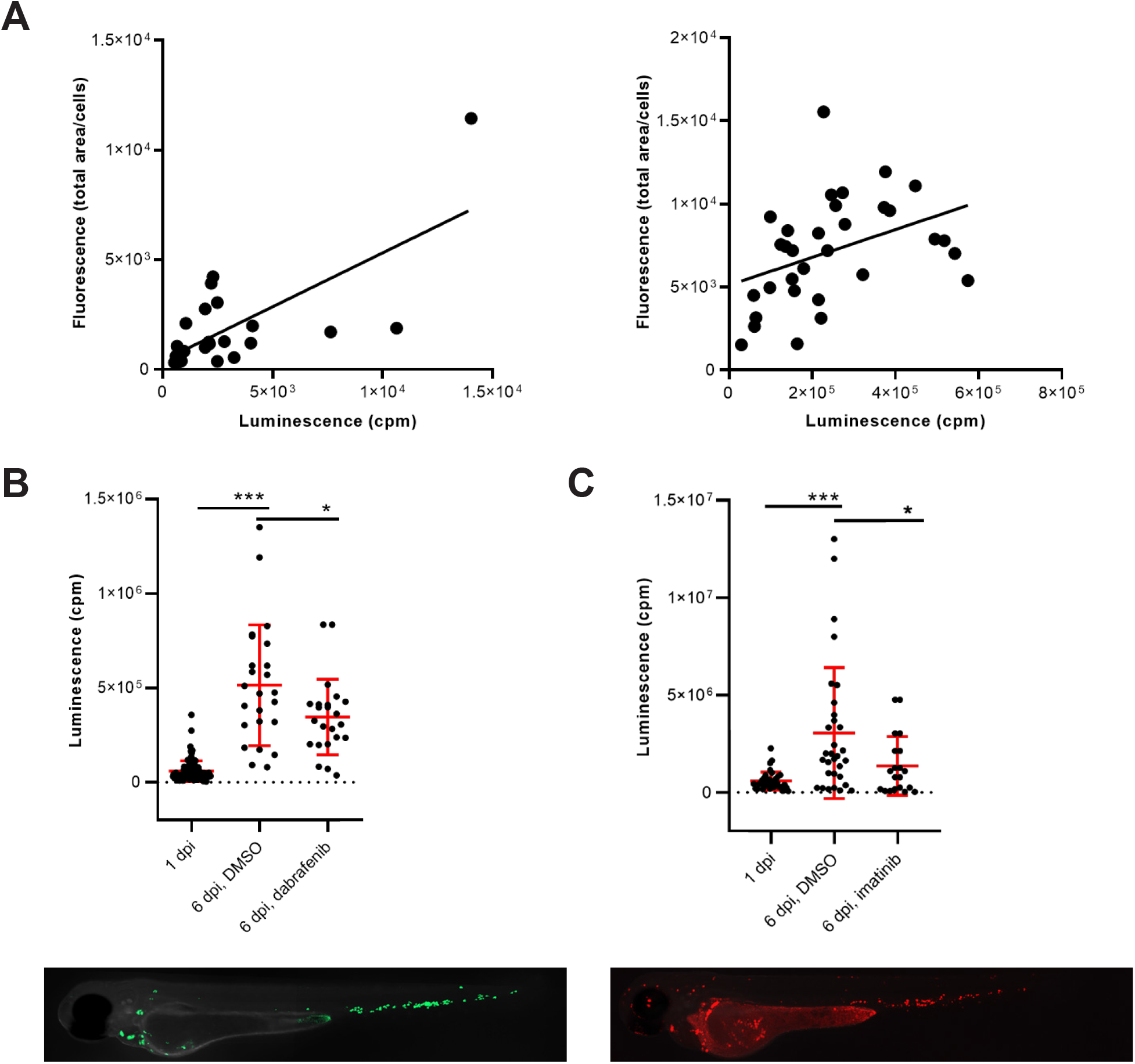
Transplanted cancer cells survive *in vivo* in zebrafish embryos and their growth can be inhibited by small molecules. (A) Correlation of fluorescence to luminescence as measured in transplanted zebrafish embryos at 1 day post-injection (dpi). Every dot is a readout from a single embryo. On the left, the correlation of EGFP to NanoLuc in ZMEL1 is shown (slope is significantly non-zero, P = 0.0001, Goodness of fit (R) = 0.4916). On the right, the correlation of mCherry to NanoLuc in K562 is shown (slope significantly non-zero, P = 0.0251, Goodness of fit (R) = 0.1563). (B) Growth and drug inhibition of growth of ZMEL1 cells in zebrafish embryos, 1 dpi - 6 dpi. ZMEL1 cells grew significantly *in vivo* from 1 dpi to 6 dpi. The growth was significantly inhibited by a BRAF inhibitor, dabrafenib. Below, there is a representative image of a 1 dpi *casper*, *prkdc*^−/−^ zebrafish embryo transplanted with green ZMEL1-*EGFP*-*NLuc* cells. (C) Growth and drug inhibition of growth of K562 cells in zebrafish embryos, 1 dpi - 6 dpi. K562 cells grew significantly *in vivo* from 1 dpi to 6 dpi. The growth was significantly inhibited by imatinib. Below, there is a representative image of a 1 dpi *casper*, *prkdc*^−/−^ zebrafish embryo transplanted with red K562-*mCherry*-*NLuc* cells. (B-C) Statistical significance was determined by unpaired two-tailed t-test. *P < 0.05 ***P < 0.001. Luminescence measured *in vivo* in single embryos is represented by a single dot in dot plots. Fluorescence images were acquired on Zeiss AxioZoom.V16 with Axiocam-506 mono camera and the ZEN Blue software.

### Validation of bioluminescent platform using dabrafenib and imatinib in vivo

Next, we initiated *in vivo* proof-of-concept cancer cell treatment to show the utility of our platform for small-molecule validation and inhibitor screening. We followed cancer cell growth *in vivo* for 5 days after transplantation and were able to accurately quantify this over time(Fig. 1B-C). We kept K562 xenografted larvae at 36°C, which is a compromise temperature, where both the larvae and human cells survive and grow normally. Well-established inhibitors of cancer cell growth were used in both zebrafish ZMEL1 allografts and K562 xenografts previously (Kansler et al., 2017; Pruvot et al., 2011). We were able to inhibit ZMEL1 cell growth with the use of 4 μM dabrafenib (BRAF^V600E^ inhibitor) *in vivo* (Fig. 1B). To inhibit K562 cell growth in zebrafish we successfully used 10 μM imatinib mesylate, a well-established kinase inhibitor of BCR-ABL as the main target (Fig. 1C). After validating the bioluminescence readout *in vivo*, we proceeded towards testing small-molecule inhibitors in a higher throughput mode. For this purpose, we selected a set of 180 known kinase inhibitors which are biologically active, target a broad spectrum of kinases, and some of which are used as drugs for the treatment of various cancer types (Table S1).

### Small-molecule screening setup and workflow

All inhibitors were first tested for *in vivo* toxicity, with 36% exhibiting toxicity at the 10 μM screening concentration. The workflow of our *in vivo* kinase inhibitor screening started with cancer cell transplantation into 2 dpf embryos. Embryos were sorted for correct transplantation of a sufficient amount of cancer cells in the blood vessels according to their fluorescent signal. Embryos were thoroughly washed, anesthetized and arranged one-by-one into wells of a 96-well plate at 1-day post-injection (1 dpi). We used uninjected embryos as a negative control to measure background luminescence. After the luciferase assay, the embryos were washed to remove all of the substrate and anesthetic and they were divided into groups of 6 into wells of a 24-well plate. In this setup, the embryos were treated with inhibitors which proved to be non-toxic from our preselected set for 5 days (1 dpi – 6 dpi). As a negative control, DMSO was used as it was the solvent of all our compounds. We used the previously validated inhibitors from our proof-of-concept experiments – imatinib for K562 and dabrafenib for ZMEL1 – as positive controls. The fresh medium and compounds were exchanged at 4 dpi and the experiment was terminated at 6 dpi (Fig. 2).

**Fig. 2.**
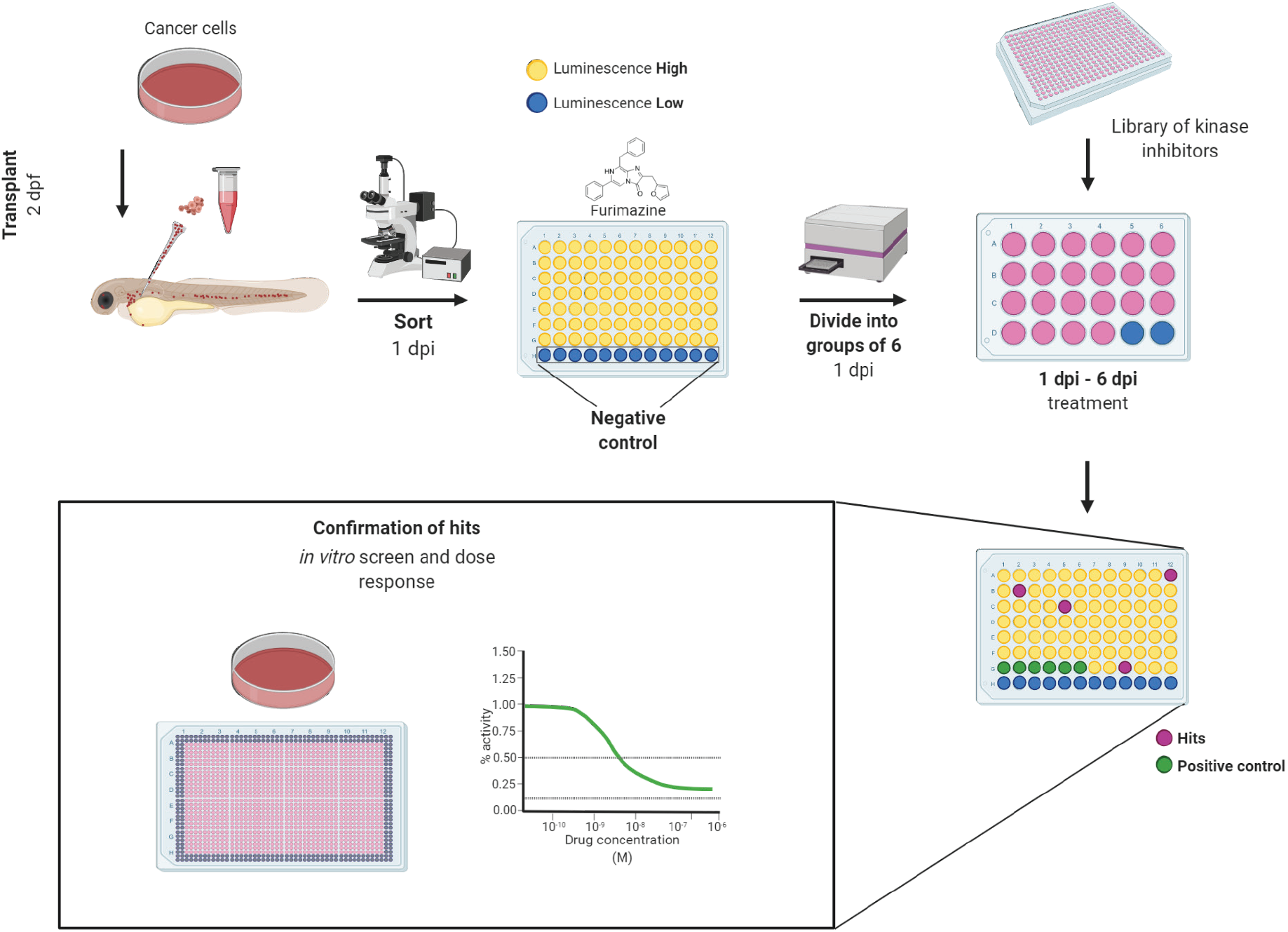
Workflow of *in vivo* small-molecule using bioluminescence screening platform. The workflow of *in vivo* small-molecule screening started with cancer cell transplantation into 2 days post-fertilization (dpf) zebrafish embryos. The transplanted embryos were washed and kept in incubator overnight. At day 1 post-injection (dpi) the embryos were sorted according to fluorescence under an Olympus macroscope and were divided into wells of a 96-well solid white plate. Uninjected embryos in E3 water and E3 water without embryos were used as negative controls to determine the background level of luminescence. Luciferase assay was carried out using the Furimazine substrate by adding it into all wells equally and the luminescence was measured after 10 minutes of incubation on the EnVision plate reader. After the measurement, embryos were recovered, washed and randomly divided into groups of 6 animals into 24-well polystyrene plates, where they were treated by inhibitors from a library of kinase inhibitors at the final concentration of 10 μM and DMSO was used as negative control. We used dabrafenib and imatinib as positive controls. At the end of the experiment, at 6 dpi, the luminescence of whole embryos was measured again to determine the cell growth or its inhibition. The lower the final luminescence, compared to positive controls treated with DMSO, the stronger is the inhibitory effect. Finally, the compounds can be analyzed *in vitro* to determine dose response curves and to compare the *in vitro* vs. *in vivo* effectivity. Figure created in bioRENDER.

Further, we re-tested a small group of inhibitors, which were moderately toxic *in vivo*, at a reduced concentration, 1 μM. We selected 10 compounds each for both ZMEL1 and K562 cells. After the *in vivo* screen, we decided to test all of the non-toxic kinase inhibitors in dose response *in vitro*. We noticed that the efficiency of inhibitors did not always correspond to their inhibitory *in vivo* effect, as shown by IC50 values from the *in vitro* screen (Table S1). This applied for around half of the active compounds found to inhibit ZMEL1 as well as K562 cell growth *in vivo*.

### Melanoma cell growth is inhibited mostly by compounds targeting cell proliferation and cell cycle

Targeting the members of the RAS-RAF-MEK-ERK signaling pathway has been shown to be beneficial in fighting various types of cancer (Germann et al., 2017; Chappell et al., 2011). Inhibition of BRAF and MEK are among the most successful current treatment strategies for fighting melanoma (Patton et al., 2021a). In our *in vivo* inhibitor screen, we found a total of 26 hit compounds (used either at 10 μM or 1 μM) which significantly inhibited the growth of melanoma cells. Most of them are known to target the members of RAS or p38 MAPK signaling pathways, for example, doramapimod, PLX-4720, cobimetinib, or SL-327 (11 compounds, Fig. 3A). Further, we found inhibitors affecting cell cycle control, mainly CDKs, to be effective against melanoma as well. These inhibitors can cause either cell cycle arrest or apoptosis, for example, abemaciclib, AMG-900, or PHA-793887 (8 compounds, Fig. 3B). The rest of the inhibitors which we found effective *in vivo*, targeted various types of receptor tyrosine kinases (RTKs, 7 compounds), for example, pazopanib, PP-121, and SB-431542 (Fig. 3C). The predicted main targets for all of our active compounds are listed in a table (Fig. 3D) according to the Probes & Drugs portal (Skuta et al., 2017). The expression of respective target genes in ZMEL1 was extracted from RNA-sequencing data (GEO accession number GSE151677) (Kansler et al., 2017) and was further normalized and analyzed using DeSeq2 (Table S2) (Love et al., 2014). All of the active compounds targeted at least one moderately expressed kinase. From all targeted kinases in melanoma, the reoccurring and supposedly most relevant ones were *mapk14a/b* and *mtor*. We composed all of this information into a comprehensive table with the expression level of each predicted target gene individually visualized by color heatmap (Fig. 3D). For additional and detailed information also see Table S1. The original expression data for each of the predicted target genes is in Table S2.

**Fig. 3.**
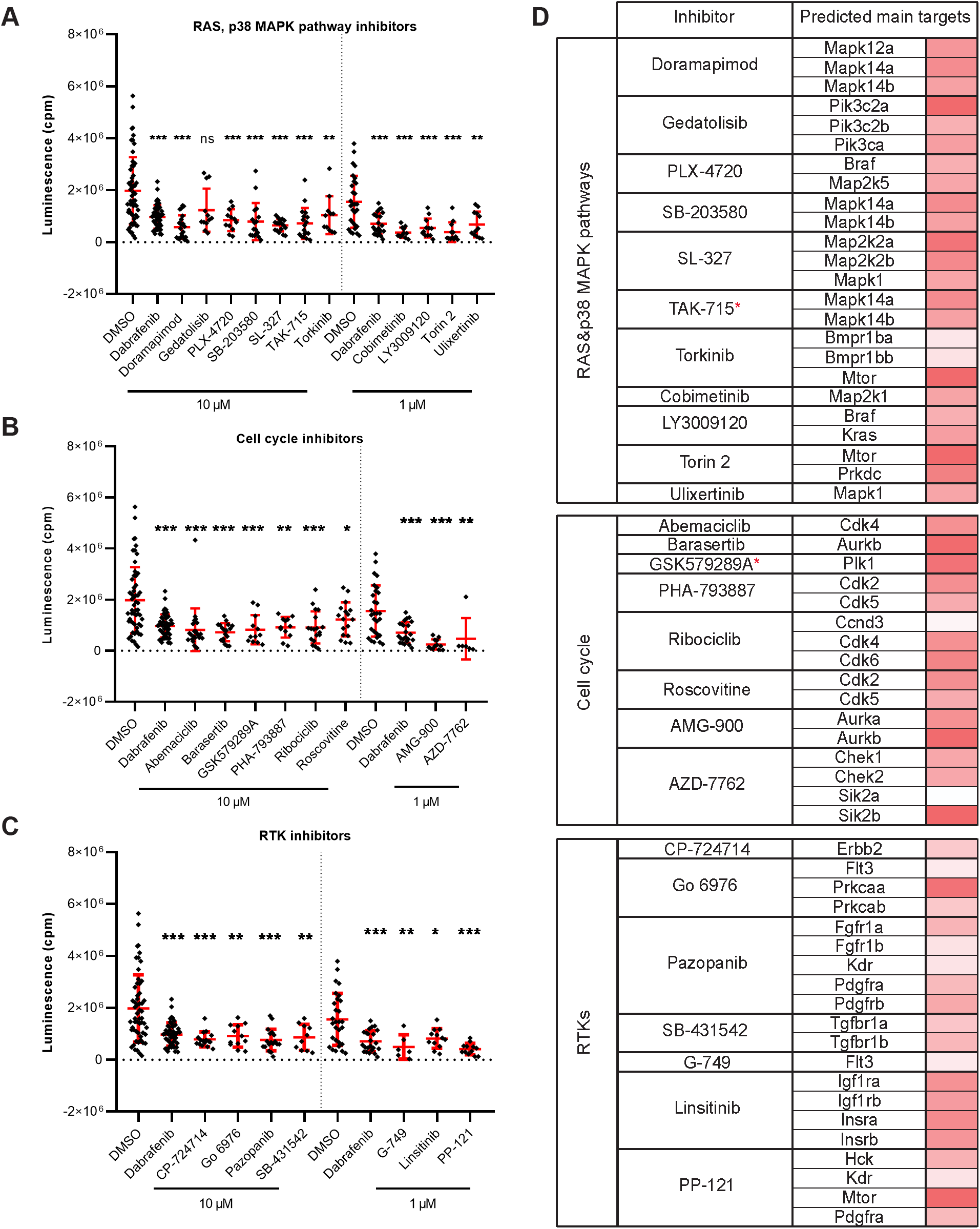
Kinase inhibitors active in transplanted ZMEL1 melanoma cells. (A) Inhibitors that targeted the RAS and p38 MAPK pathways and significantly inhibited melanoma cell growth *in vivo*. (B) Inhibitors that targeted cell cycle related proteins and significantly inhibited melanoma cell growth *in vivo*. (C) Inhibitors that targeted receptor tyrosine kinases (RTKs) and significantly inhibited melanoma cell growth *in vivo*. (A-C) Statistical significance was determined by Mann-Whitney test. *P < 0.04, **P <0.002, ***P < 0.001. Luminescence measured *in vivo* in single embryos is represented by a single dot in dot plots. All experiments were done in 2 – 3 repeats. (D) List of all the inhibitors from A-C with their predicted main protein targets in zebrafish ZMEL1 cells. The last column with red bars on the right represents the average expression of individual target genes in zebrafish cells which was extracted from a publicly available RNA-sequencing dataset. Inhibitors with a potent new function in melanoma are labeled with a red asterisk.

### Leukemic cell growth is inhibited by targeting cell proliferation, migration and survival

Leukemia is commonly induced by a combined effect of multiple genetic alterations which can hinder the establishment of targeted therapy (Vetrie et al., 2020). Targeting the BCR-ABL1 fusion gene, growth factor receptors (GFRs) and also the RAS-RAF-MEK-ERK pathway, all belong to strategies used in the treatment of myeloid leukemias (Bhullar et al., 2018). We found 17 compounds (used again either at 10 μM or 1 μM) as hits in our screen looking at the inhibition of leukemia cancer cell growth *in vivo*. Out of these, 9 compounds, for example, NG-25, or ipatasertib, are predicted to target members of the RAS or p38 MAPK signaling pathways. We also showed the strong effect of GNF-5, a selective BCR-ABL1 inhibitor, *in vivo* (Fig. 4A). Another 8 compounds, for example, BAY-826, AZD-7762, or DDR-IN-1 target cell migration and cell cycle-related protein kinases (Fig. 4B). In a table (Fig. 4C), we listed the predicted main targets for all of our hit compounds together with the target gene expression in the K562 cell line. In leukemic cells, we have found active compounds targeted at moderately to highly expressed kinases. From all targeted kinases in leukemia, the reoccurring and supposedly most relevant ones were *MAPK14, DDR1* and *AURKA*. The latter was extracted from an RNA-seq dataset (GEO accession PRJNA30709) and further analyzed similarly as in the case of the ZMEL1 cell line (Table S2).

**Fig. 4.**
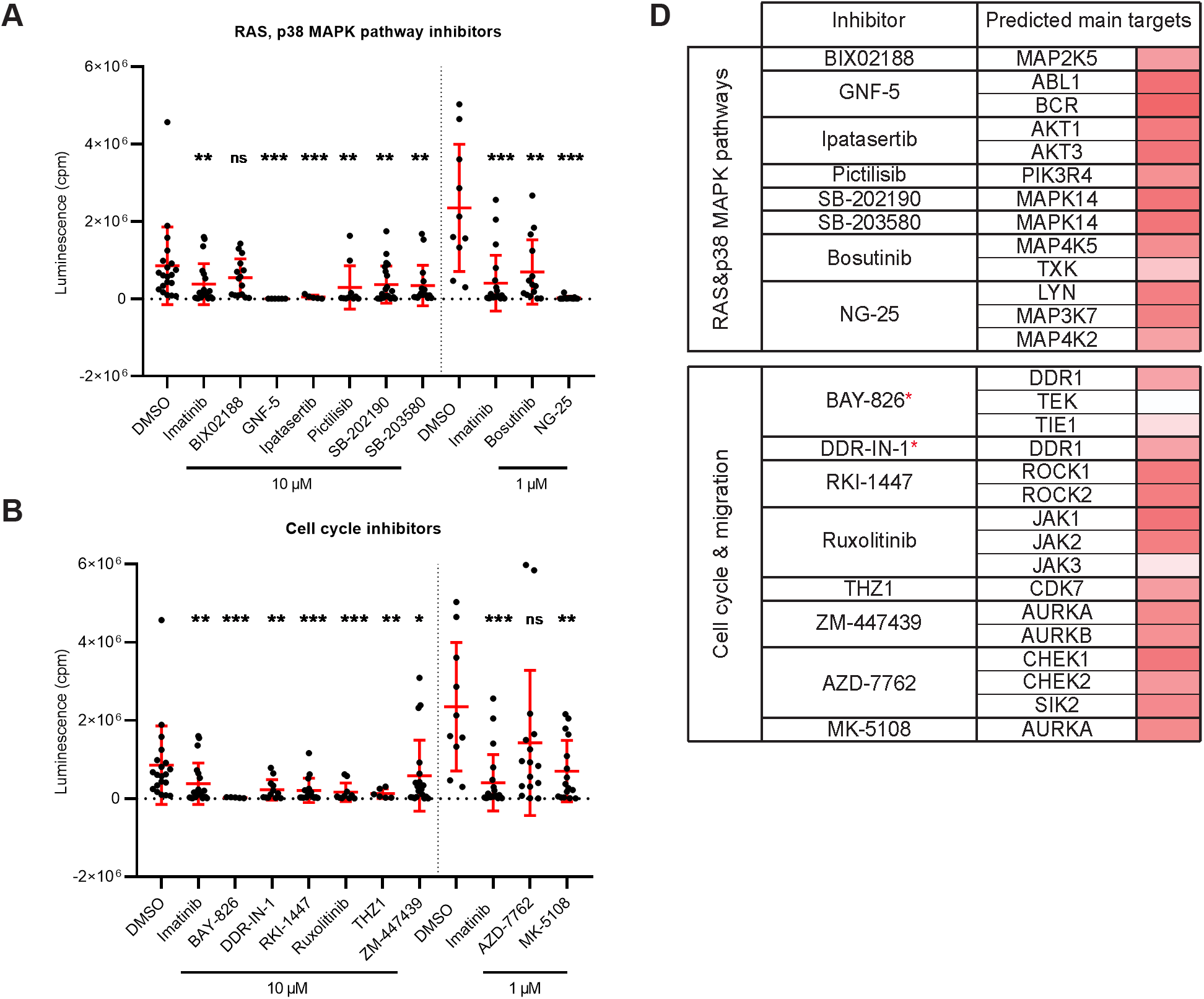
Kinase inhibitor active in transplanted leukemia K562 cells. (A) Inhibitors that targeted the RAS and p38 MAPK pathways and significantly inhibited leukemia cell growth *in vivo*. (B) Inhibitors that targeted cell cycle and cell migration related proteins and significantly inhibited leukemia cell growth *in vivo*. (A-B) Statistical significance was determined by Mann-Whitney test. *P < 0.04, **P <0.002, ***P < 0.001. Luminescence measured *in vivo* in single embryos is represented by a single dot in dot plots. All experiments were done in 2 – 3 repeats. (C) List of all the inhibitors from A-B with their predicted main protein targets in human K562 cells. The last column with red bars on the right represents the average expression of individual target genes in human cells which was extracted from a publicly available RNA-sequencing dataset. Inhibitors with a potent new function in leukemia are labeled with a red asterisk.

To summarize, we found multiple small molecules predicted or known to target various signaling pathways in our *in vivo* zebrafish screen which inhibit melanoma (Fig. 5A) and CML cell growth (Fig. 5B). Our results provide proof of concept that zebrafish can be utilized as a reliable model for cancer drug discovery in a medium/high-throughput setup with bioluminescence as a readout. This model would enable fast and efficient assessment of drug toxicity and efficacy *in vivo* and could serve as a platform for drug repurposing experiments.

**Fig. 5.**
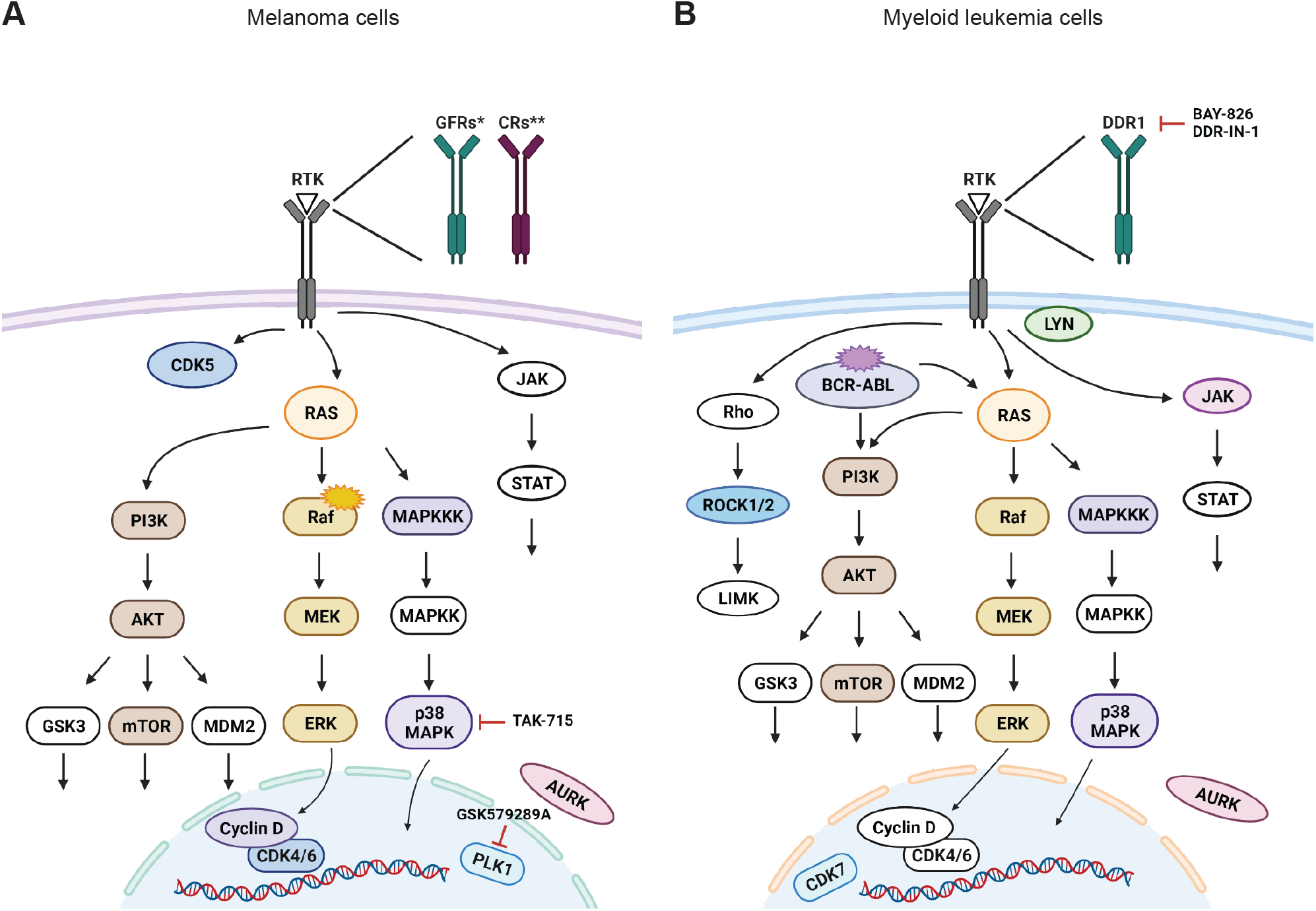
Targeted signaling pathways as predicted from *in vivo* kinase inhibitor screen. For simplicity, we show the predicted human proteins in this figure without the zebrafish paralogs. Proteins, that were targeted by our kinase inhibitor set are depicted in color, non-targeted ones are white. Selected inhibitors with a potent new function are depicted in the schemes as well. Signaling pathways in (A) BRAF^V600E^ mutant melanoma cells and in (B) cells of chronic myelogenous leukemia (CML) with BCR-ABL translocation. Figure created in bioRENDER.

## DISCUSSION

Human tumor cell behavior is commonly modeled in clinically relevant settings relying on *in vitro* organoids or *in vivo* mouse xenografts. Recently, zebrafish embryos were shown to be routinely transplanted by cancer cells and further used for the evaluation of tumor growth and dissemination (Kirchberger et al., 2017; Konantz et al., 2012; Potts and Bowman, 2017; Pruitt et al., 2017). Grafted embryos and larvae could be treated by simply soaking them in chemical compounds dissolved in water. Cell proliferation is most often measured until 3-4 dpi by the quantification of fluorescently labeled cancer cells, *in vivo* (Corkery et al., 2011; Ghotra et al., 2012; Hason and Bartunek, 2019; Patton et al., 2021b; Tulotta et al., 2016; Tulotta et al., 2019) or *ex vivo* (Somasagara et al., 2021). Our screening workflow enables rapid characterization of tumor-inhibitor responsiveness within a time frame of seven days.

Luminescence can be measured multiple times during the experiment as the procedure is noninvasive and the NanoLuc substrate is not toxic for animals. The simplicity of the luciferase assay allowed us to screen hundreds of zebrafish larvae within minutes. Therefore, this approach could be utilized for medium- to high-throughput pre-clinical screening of chemical compounds targeting any cancer type transplanted into zebrafish embryos. Further, it offers a unique opportunity to rapidly validate and characterize the inhibitory effect of drugs individually or in combinations. Our zebrafish xenograft model could be a platform of choice to accelerate drug discovery and repurposing.

Our first aim was to validate bioluminescence in a zebrafish model of cancer cell growth. Bioluminescent imaging has been previously used in zebrafish to visualize hematopoietic cell proliferation, tumorigenesis or apoptosis *in vivo* (Astuti et al., 2017; de Latouliere et al., 2021; Manni et al., 2019; Tobia et al., 2021). We showed that dabrafenib reduced melanoma growth and imatinib reduced CML growth in transplanted zebrafish larvae by measuring luminescence. Both compounds were used in earlier studies with similar results (Ablain et al., 2021; Cohen et al., 2021; Corkery et al., 2011; Pruvot et al., 2011). After we validated our experimental setup, we continued with the *in vivo* screen of kinase inhibitors. Additionally, we tested a small group of inhibitors that were on the borderline of toxicity, at 10 times lower final concentration. It can be difficult to set up a correct therapeutic dose for inhibitors targeting a vast number of targets, even if they belong to the same family of target proteins, as kinases. Usually, a trade-off average concentration has to be selected within the activity range of all small-molecule compounds used. This value is generally within the range of 1 – 15 μM in most zebrafish *in vivo* pharmaceutical screens (Bowman and Zon, 2010; Colanesi et al., 2012; Haney et al., 2021; Oprisoreanu et al., 2021; Precazzini et al., 2020). Therefore, we decided to use 10 μM as our selected *in vivo* screening concentration. Unfortunately, due to the complexity of the screening protocol, it is not always feasible to test more than one inhibitor concentration without losing throughput. Additionally, most drugs work similarly in zebrafish and humans, although the affinity of a drug for the human protein may differ from its affinity for the zebrafish protein. This might pose a problem, and as a result, different dosing might be required in mammals (Cagan et al., 2019; Fazio et al., 2020; Yan et al., 2020).

Our *in vivo* data suggested that the most affected cell processes, for both melanoma and CML, are MAPK signaling and cell cycle-related signaling (Fig. 5A, 5B). The RAS-RAF-MEK-ERK signaling pathway in particular is an attractive therapeutic target in numerous cancer types (Chappell et al., 2011). In melanoma, especially BRAF and MEK-targeted therapy was proved to be beneficial (Flaherty et al., 2012; Patton et al., 2021a). Further, BRAF is mutated in approximately 50% of cutaneous melanoma cases in human patients (Ribas et al., 2019; Robert et al., 2019). In our experiments, we used the melanoma cell line ZMEL1, which carries human *BRAF*^V600E^, the most frequent mutant variant found in patients (Heilmann et al., 2015; Patton et al., 2021a; Patton et al., 2005). From this group of hit compounds, we identified TAK-715 as a promising small molecule actively inhibiting melanoma growth *in vivo*. This inhibitor was previously characterized as a pharmaceutical candidate for use in rheumatoid arthritis (Miwatashi et al., 2005). Cell cycle affecting inhibitors was the second most represented type of compounds reducing melanoma growth in our experiments. From all of them, GSK579289A may be a promising new inhibitor targeting polo-like kinase 1 (PLK1) (Rheault et al., 2010). PLK1 was shown to be overexpressed in melanoma (Takai et al., 2005) and its therapeutic targeting could be beneficial in various types of cancer (Gutteridge et al., 2016). Activating mutations of RTKs can lead to abnormal downstream signaling, aberrant growth and survival of malignant melanocytes. RTKs have been connected to melanoma pathogenesis and are considered potent therapeutic targets (Easty et al., 2011; Sabbah et al., 2021). In our screen, we found various RTK inhibitors efficiently inhibiting ZMEL cell growth *in vivo*.

Similarly, in myeloid leukemia, inhibitors targeting MAPK signaling pathways were shown previously to be beneficial as mono- as well as combination therapy options (Rocca et al., 2018; Tambe et al., 2020; Yen et al., 2020). Additionally, we demonstrated that the inhibition of proteins related to cell cycle and migration can be useful in targeting leukemic expansion *in vivo*. Inhibitors targeting discoidin domain receptor 1 (DDR1), BAY-826 and DDR-IN-1 are promising hits as DDR1 inhibition has potential in cancer therapy (Berestjuk et al., 2022; Elkamhawy et al., 2021). Moreover, DDR1 therapeutic targeting *in vivo* was not yet characterized for the treatment of myeloid leukemia.

The inhibition efficiency *in vitro* did not fully correspond to the inhibitory effects observed *in vivo*, which likely relates to pharmacokinetic and/or pharmacodynamic properties. This is an advantage of the zebrafish *in vivo* screen, as it provides a full picture of cancer growth inhibition as it is not fully recapitulated *in vitro*. Indeed, it is known that *in vivo* assays can detect inhibitors that act in a non-cell autonomous manner in the animal, therefore efficiently reducing tumor progression even when inactive *in vitro* (Sonoshita et al., 2018). In addition, polypharmacology can support the overall efficacy of a certain chemical compound by targeting not only targets in tumor cells but also cells in the tumor microenvironment (Cagan et al., 2019; Dar et al., 2012; Edwards et al., 2019).

Overall, our study demonstrates the utility of the zebrafish platform employing bioluminescence as a readout for fast and efficient drug screening. The usefulness of this model could be employed not only in drug repurposing but also in *de novo* drug discovery.

## MATERIALS AND METHODS

### Animals

Zebrafish were kept and raised in ZebTEC aquatic systems (Tecniplast) according to standard procedures (Alestrom et al., 2020) and were tracked using Zebrabase (Oltova et al., 2018). The immunodeficient *prkdc*^fb103/fb103^ zebrafish strain (Moore et al., 2016), herein referred to as *prkdc*^−/−^, was used as transplantation recipients. Wild-type (AB) or *casper* strains (White et al., 2008) were used as controls. Zebrafish embryos were dechorionated using pronase (Roche) at 24 hours post fertilization (hpf) and were kept in E3 medium (Westerfield, 2007) up to the larval stage. Procedures for animal husbandry and experimentation were approved by the Animal Ethics Committee of the Institute of Molecular Genetics (13/2016 and 96/2018) in compliance with national and institutional guidelines.

### Cell culture and preparation of cancer cells with double reporter system

K562 (ATCC) human erythroleukemia cells were grown in Iscove’s Modified Dulbecco’s Medium (IMDM, Gibco) supplemented with 1% Glutamax (Gibco), 10% fetal bovine serum (FBS, Gibco), and 1% Penicillin-Streptomycin (P-S, Gibco) at 37°C/5% CO_2_ and were split every 3 days at 1:10 ratio. ZMEL1 zebrafish melanoma cells were grown in high glucose Dulbecco’s Modified Eagle’s Medium (DMEM, Gibco) medium supplemented with 1% Glutamax, 10% FBS, and 1% P-S and were split approximately every 4 days at a 1:4 ratio. ZMEL1 cells were kept in the incubator at 28.5°C/5% CO_2_.

A non-secreted version of the luciferase enzyme, NanoLuc^®^ (NLuc, Promega), was cloned into a bicistronic lentiviral expression vector pLVX-EF1α-IRES-mCherry (Clontech) creating pLVX-EF1α-IRES-mCherry-NLuc. This vector allows the simultaneous coexpression of *mCherry* and *NLuc*.

To prepare K562-*mCherry*-*NLuc* cells, semiconfluent HEK293FT cells were first co-transfected by Lipofectamine 2000 (Invitrogen) for the production of lentiviral particles; a mixture of *pLVX-EF1α-IRES-mCherry-NLuc*, *pVSV-G* (Addgene #8454) and *psPAX2* (Addgene #12260) was prepared in 3:1:2 ratio and the cells were transduced as described previously (Konirova et al., 2017). After 48 hours the viral supernatant was harvested and centrifuged. Lentiviral particles were precipitated by PEG-it (System Biosciences) as described by the manufacturer and were resuspended in sterile 1xPBS and frozen in aliquots at −80°C. K562 cells were incubated with lentiviral particles for 24 hours and then the medium was exchanged. Cells were single-cell sorted for *mCherry*+ by BD Influx cell sorter into 96-well plates at 48 – 72 hours post infection. Single clones were cultivated and used in further experiments.

ZMEL1-*EGFP* cells were isolated from a transgenic *mitfa-BRAF*^V600E^*;mitfa-EGFP;p53*^−/−^ fish as previously described (Heilmann et al., 2015). A tumor was excised and a stable cell line was derived, which was *EGFP* labelled due to *mitfa* promoter expression. To create the ZMEL1*-EGFP-NanoLuc* line, a plasmid was created in which the *ubb* promoter (Mosimann et al., 2011) was used to drive the luciferase open reading frame, followed by an SV40 polyadenylation signal. This was cloned into a Tol2-based vector backbone that contains a blasticidin selection cassette. The plasmid was electroporated into the ZMEL1-*EGFP* line using the Neon electroporation system. Following a 24-hour recovery for cells to attach, they were placed under blasticidin selection for 2 weeks. After 2 weeks, the cells were returned to normal media and were single-cell sorted from a mass culture for *EGFP*+ by BD Influx cell sorter into 96-well plates. Single clones were grown to confluency in DMEM/15% FBS and were used in transplantation experiments.

### Luciferase assay, *in vitro* and *in vivo*

Furimazine (Promega), a substrate of NanoLuc, was used according to the manufacturer protocol in *in vitro* screens. Briefly, cells were seeded into 96-well solid white plates (Corning) 24 hours prior the luciferase assay. Full media without phenol red was used. Prior to measurement, all plates were allowed to reach ambient temperature and the NanoGlo reagent with Furimazine (Promega) was added in 1:1 ratio to the cells. Cells were briefly shaken, incubated for 10 minutes, and the luminescence was measured on an EnVision plate reader (Perkin Elmer).

For luminescence measurements *in vivo*, 50x diluted Furimazine was used, diluted in 5% ethanol instead of the NanoGlo lysis reagent. Zebrafish embryos and larvae were measured at 1 day post injection (dpi) and 6 dpi. They were thoroughly washed in E3 medium, anesthetized by 1x Tricaine (ethyl 3-aminobenzoate methanesulfonate/MS-222, 0.16 mg/mL) and distributed into wells of a 96-well solid white plate (Corning) in approximately 50 μL of E3/1x Tricaine. Furimazine was added in a ratio 1:2 to the embryos, the plates were briefly shaken and incubated for 10 minutes at room temperature. Luminescence was measured on EnVision.

### Transplantation of cancer cells into zebrafish embryos

Cancer cells were washed in 1x PBS, filtered, counted and diluted to a final concentration of 60 x 10^6^ cells/ml in 2% PVP40 (polyvinylpyrrolidone, Sigma; PVP was diluted in full DMEM). Phenol red (Sigma) was added to the mixture to better visualize the injected cells in real time.

At 2 days post fertilization (dpf) embryos were anesthetized by 1x Tricaine and laterally arranged in groups of 25 – 50 on a 2% agarose dish. Borosilicate glass capillaries (Harvard Apparatus, GC100FS-10) were pulled to create microinjection needles which were then opened by tweezers (≈20 μm). Cancer cell mixture was filled into the capillary and ≈100 cells were transplanted to the blood stream of embryos through the dorsal part of the duct of Cuvier. After transplantation, embryos were recovered and washed in E3 and incubated at 35°C/K562 or 28°C/ZMEL1 until the next morning when they were sorted according to fluorescence. All embryos with an insufficient number of transplanted cells were discarded at this point.

### Selection of kinase inhibitor set and in vivo inhibitor screen

A set of 180 kinase inhibitors was selected based on data from the Probes & Drugs portal (Skuta et al., 2017). For each kinase, a maximum of 3 inhibitors was selected based on their potency, diversity and availability. Compounds labelled as chemical probes were prioritized, while compounds labelled as obsolete/historic (Arrowsmith et al., 2015) were removed from the library.

The toxicity of all inhibitors at 10 μM concentration was first tested in our setup, and then for subsequent treatment experiments and the *in vitro* inhibitor screen, only the non-toxic inhibitors were used. For the *in vivo* screen of inhibitors, zebrafish embryos transplanted with either ZMEL1 or K562 cells, were sorted according to cancer cell fluorescence. At 1 dpi the sorted embryos were washed, anesthetized, aligned into wells of white opaque 96-well plates (Corning) for luminescence measurement. Embryos were then washed in fresh E3 medium and kept in groups of 6 in 24-well polystyrene plates (Nunc) in 1 ml of E3 water. Embryos were treated by a final 10 μM dose of each non-toxic inhibitor and 1 μM dose for a selected group of borderline toxic inhibitors. Positive controls were imatinib mesylate (10 μM) to treat K562 transplanted larvae and dabrafenib (4 μM) for ZMEL1 transplanted larvae. DMSO (0.1%) was used as a negative control. The experiment was terminated at 6 dpi after the final luminescence measurement, with larvae being euthanized by Tricaine overdose on ice.

### *In vitro* inhibitor dose response and data evaluation

K562-*mCherry*-*NLuc* cells were washed in 1x PBS, filtered using a 40 μm strainer, counted and seeded using a reagent dispenser Tempest (Formulatrix) at the density of 2.5 x 10^3^ cells/ml into 1536-well solid white plates (Corning). Rim wells were filled with IMDM medium to reduce edge effect. Inhibitors were then dispensed by an acoustic liquid handler Echo 525 (Beckman Coulter). Compounds were tested in 12 concentration points extending from 100 μM to 0.1 nM, in triplicates. DMSO was used as the negative control and imatinib as the positive control. Wells with medium and DMSO backfill were used to measure background luminescence. The plates with added compounds were shaken, briefly spun down and put back to the incubator for 3 days. NanoGlo reagent was dispensed by Tempest in the ratio 1:1, the cells were shaken, spun down and luminescence was measured using a PHERAstar FSX microplate reader (BMG Labtech). Signal intensity is proportional to the number of live cells in the sample. Data were collected, normalized, and processed using proprietary LIMS system ScreenX.

For *ZMEL1*-*EGFP*-*NLuc* the workflow was the same but for the following exceptions. Cells were seeded at a density of 2 x 10^5^ cells/ml. All the plates with cells were then put into wet chambers and were kept in an incubator until the next day to let the cells attach. Dabrafenib was used as the positive control. NanoGlo reagent was dispensed in the ratio 1:2.5 by Tempest.

### RNA-seq data analysis

RNA-seq datasets for the ZMEL1 cell line were downloaded from GEO (Accession number GEO: GSE151677). The quality of reads was checked with FastQC (de Sena Brandine and Smith, 2019) and MultiQC tools (Ewels et al., 2016). Mapping to the reference *Danio rerio* transcriptome (Ensembl, release 104) was performed with Salmon (Patro et al., 2017). The obtained count matrix was imported in R, and analyzed with DESeq2 (Love et al., 2014). The reads were normalized using TPM and plotted into box plots using the Tidyverse R package (Wickham et al., 2019).

Datasets for the K562 cell line were downloaded from GEO (Accession number GEO: PRJNA30709 and PRJEB7858) and the data analysis was the same as for ZMEL1 with the exception that mapping was performed to the reference *Homo sapiens* transcriptome (Ensembl, release 104).

### Fluorescence imaging and quantification

Fluorescent imaging was done for correlation of fluorescence to luminescence experiments at 3 dpf in both K562 and ZMEL1 transplanted embryos. Images were taken in multiple z-stacks using Zeiss AxioZoom.V16 with Axiocam 506 mono camera. The magnification was 80x. Orthogonal projections were created in ZEN Blue 2.3 software. The area of fluorescent cells in the whole embryos was calculated in Fiji.(Schindelin et al., 2012)

### Statistical analysis

P values for *in vivo* experiments were calculated by the unpaired two-tailed Student’s *t*-test or the nonparametric Mann-Whitney U test using GraphPad Prism 8.0.1.

The correlation of fluorescence to luminescence signal was done in GraphPad Prism 8.0.1 by fitting the two variables into simple linear regression.

For the *in vitro* inhibitor dose response and to calculate IC50 values GraphPad Prism 8.0.1 was used.

## Acknowledgements

We would like to thank Nikol Pavlu, Tereza Hojerova and Tereza Hingarova for animal care; Ctibor Skuta for his help with the selection of the kinase inhibitor set, Martin Popr for preparing the chemical library for experiments, Trevor Epp for reading and editing the manuscript. We thank David M. Langenau for providing the *prkdc*^D3612fs^ mutant fish line.

## Author contributions

M.H and P.B. designed the research, M.H., J.J., I.V., T.S.-V. performed the experiments. M.H., J.J., I.V., M.B., R.M.W. and P.B. analyzed the data, M.H., J.J. and P.B. wrote the manuscript.

## Supplementary information

Supplementary information available online at …

## References

Ablain, J., Liu, S., Moriceau, G., Lo, R. S. and Zon, L. I. (2021). SPRED1 deletion confers resistance to MAPK inhibition in melanoma. J Exp Med 218.

Alestrom, P., D’Angelo, L., Midtlyng, P. J., Schorderet, D. F., Schulte-Merker, S., Sohm, F. and Warner, S. (2020). Zebrafish: Housing and husbandry recommendations. Lab Anim 54, 213–224.

Almstedt, E., Rosén, E., Gloger, M., Stockgard, R., Hekmati, N., Koltowska, K., Krona, C. and Nelander, S. (2021). Real-time evaluation of glioblastoma growth in patient-specific zebrafish xenografts. Neuro-Oncology.

Arrowsmith, C. H., Audia, J. E., Austin, C., Baell, J., Bennett, J., Blagg, J., Bountra, C., Brennan, P. E., Brown, P. J., Bunnage, M. E. et al. (2015). The promise and peril of chemical probes. Nat Chem Biol 11, 536–41.

Astuti, Y., Kramer, A. C., Blake, A. L., Blazar, B. R., Tolar, J., Taisto, M. E. and Lund, T. C. (2017). A Functional Bioluminescent Zebrafish Screen for Enhancing Hematopoietic Cell Homing. Stem Cell Reports 8, 177–190.

Baeten, J. T., Waarts, M. R., Pruitt, M. M., Chan, W. C., Andrade, J. and de Jong, J. L. O. (2019). The side population enriches for leukemia-propagating cell activity and Wnt pathway expression in zebrafish acute lymphoblastic leukemia. Haematologica 104, 1388–1395.

Berestjuk, I., Lecacheur, M., Carminati, A., Diazzi, S., Rovera, C., Prod’homme, V., Ohanna, M., Popovic, A., Mallavialle, A., Larbret, F. et al. (2022). Targeting Discoidin Domain Receptors DDR1 and DDR2 overcomes matrix-mediated tumor cell adaptation and tolerance to BRAF-targeted therapy in melanoma. EMBO Mol Med 14, e11814.

Bhullar, K. S., Lagaron, N. O., McGowan, E. M., Parmar, I., Jha, A., Hubbard, B. P. and Rupasinghe, H. P. V. (2018). Kinase-targeted cancer therapies: progress, challenges and future directions. Mol Cancer 17, 48.

Bowman, T. V. and Zon, L. I. (2010). Swimming into the future of drug discovery: in vivo chemical screens in zebrafish. ACS Chem Biol 5, 159–61.

Cagan, R. L., Zon, L. I. and White, R. M. (2019). Modeling Cancer with Flies and Fish. Dev Cell 49, 317–324.

Capasso, A., Lang, J., Pitts, T. M., Jordan, K. R., Lieu, C. H., Davis, S. L., Diamond, J. R., Kopetz, S., Barbee, J., Peterson, J. et al. (2019). Characterization of immune responses to anti-PD-1 mono and combination immunotherapy in hematopoietic humanized mice implanted with tumor xenografts. J Immunother Cancer 7, 37.

Cohen, P., Cross, D. and Janne, P. A. (2021). Kinase drug discovery 20 years after imatinib: progress and future directions. Nat Rev Drug Discov 20, 551–569.

Colanesi, S., Taylor, K. L., Temperley, N. D., Lundegaard, P. R., Liu, D., North, T. E., Ishizaki, H., Kelsh, R. N. and Patton, E. E. (2012). Small molecule screening identifies targetable zebrafish pigmentation pathways. Pigment Cell Melanoma Res 25, 131–43.

Corkery, D. P., Dellaire, G. and Berman, J. N. (2011). Leukaemia xenotransplantation in zebrafish--chemotherapy response assay in vivo. Br J Haematol 153, 786–9.

Dar, A. C., Das, T. K., Shokat, K. M. and Cagan, R. L. (2012). Chemical genetic discovery of targets and anti-targets for cancer polypharmacology. Nature 486, 80–4.

de Latouliere, L., Manni, I., Ferrari, L., Pisati, F., Totaro, M. G., Gurtner, A., Marra, E., Pacello, L., Pozzoli, O., Aurisicchio, L. et al. (2021). MITO-Luc/GFP zebrafish model to assess spatial and temporal evolution of cell proliferation in vivo. Sci Rep 11, 671.

de Sena Brandine, G. and Smith, A. D. (2019). Falco: high-speed FastQC emulation for quality control of sequencing data. F1000Res 8, 1874.

Easty, D. J., Gray, S. G., O’Byrne, K. J., O’Donnell, D. and Bennett, D. C. (2011). Receptor tyrosine kinases and their activation in melanoma. Pigment Cell Melanoma Res 24, 446–61.

Edwards, D. K.. V, Watanabe-Smith, K., Rofelty, A., Damnernsawad, A., Laderas, T., Lamble, A., Lind, E. F., Kaempf, A., Mori, M., Rosenberg, M. et al. (2019). CSF1R inhibitors exhibit antitumor activity in acute myeloid leukemia by blocking paracrine signals from support cells. Blood 133, 588–599.

Elkamhawy, A., Lu, Q., Nada, H., Woo, J., Quan, G. and Lee, K. (2021). The Journey of DDR1 and DDR2 Kinase Inhibitors as Rising Stars in the Fight Against Cancer. Int J Mol Sci 22.

Ewels, P., Magnusson, M., Lundin, S. and Kaller, M. (2016). MultiQC: summarize analysis results for multiple tools and samples in a single report. Bioinformatics 32, 3047–8.

Fazio, M., Ablain, J., Chuan, Y., Langenau, D. M. and Zon, L. I. (2020). Zebrafish patient avatars in cancer biology and precision cancer therapy. Nat Rev Cancer 20, 263–273.

Fior, R., Póvoa, V., Mendes, R. V., Carvalho, T., Gomes, A., Figueiredo, N. and Ferreira, M. G. (2017). Single-cell functional and chemosensitive profiling of combinatorial colorectal therapy in zebrafish xenografts. Proceedings of the National Academy of Sciences 114, E8234–E8243.

Flaherty, K. T., Infante, J. R., Daud, A., Gonzalez, R., Kefford, R. F., Sosman, J., Hamid, O., Schuchter, L., Cebon, J., Ibrahim, N. et al. (2012). Combined BRAF and MEK Inhibition in Melanoma with BRAF V600 Mutations. New England Journal of Medicine 367, 1694–1703.

Germann, U. A., Furey, B. F., Markland, W., Hoover, R. R., Aronov, A. M., Roix, J. J., Hale, M., Boucher, D. M., Sorrell, D. A., Martinez-Botella, G. et al. (2017). Targeting the MAPK Signaling Pathway in Cancer: Promising Preclinical Activity with the Novel Selective ERK1/2 Inhibitor BVD-523 (Ulixertinib). Mol Cancer Ther 16, 2351–2363.

Ghotra, V. P., He, S., de Bont, H., van der Ent, W., Spaink, H. P., van de Water, B., Snaar-Jagalska, B. E. and Danen, E. H. (2012). Automated whole animal bio-imaging assay for human cancer dissemination. PLoS One 7, e31281.

Gutierrez, A., Pan, L., Groen, R. W., Baleydier, F., Kentsis, A., Marineau, J., Grebliunaite, R., Kozakewich, E., Reed, C., Pflumio, F. et al. (2014). Phenothiazines induce PP2A-mediated apoptosis in T cell acute lymphoblastic leukemia. J Clin Invest 124, 644–55.

Gutteridge, R. E., Ndiaye, M. A., Liu, X. and Ahmad, N. (2016). Plk1 Inhibitors in Cancer Therapy: From Laboratory to Clinics. Mol Cancer Ther 15, 1427–35.

Hall, M. P., Unch, J., Binkowski, B. F., Valley, M. P., Butler, B. L., Wood, M. G., Otto, P., Zimmerman, K., Vidugiris, G., Machleidt, T. et al. (2012). Engineered luciferase reporter from a deep sea shrimp utilizing a novel imidazopyrazinone substrate. ACS Chem Biol 7, 1848–57.

Hanahan, D. and Weinberg, R. A. (2000). The Hallmarks of Cancer. Cell 100, 57–70.

Hanahan, D. and Weinberg, R. A. (2011). Hallmarks of cancer: the next generation. Cell 144, 646–74.

Haney, M. G., Wimsett, M., Liu, C. and Blackburn, J. S. (2021). Protocol for rapid assessment of the efficacy of novel Wnt inhibitors using zebrafish models. STAR Protoc 2, 100433.

Hason, M. and Bartunek, P. (2019). Zebrafish Models of Cancer-New Insights on Modeling Human Cancer in a Non-Mammalian Vertebrate. Genes (Basel) 10.

Heilmann, S., Ratnakumar, K., Langdon, E. M., Kansler, E. R., Kim, I. S., Campbell, N. R., Perry, E. B., McMahon, A. J., Kaufman, C. K., van Rooijen, E. et al. (2015). A Quantitative System for Studying Metastasis Using Transparent Zebrafish. Cancer Res 75, 4272–4282.

Howe, K. Clark, M. D. Torroja, C. F. Torrance, J. Berthelot, C. Muffato, M. Collins, J. E. Humphray, S. McLaren, K. Matthews, L. et al. (2013). The zebrafish reference genome sequence and its relationship to the human genome. Nature 496, 498–503.

Chappell, W. H., Steelman, L. S., Long, J. M., Kempf, R. C., Abrams, S. L., Franklin, R. A., Bäsecke, J., Stivala, F., Donia, M., Fagone, P. et al. (2011). Ras/Raf/MEK/ERK and PI3K/PTEN/Akt/mTOR inhibitors: rationale and importance to inhibiting these pathways in human health. Oncotarget 2, 135–164.

Kansler, E. R., Verma, A., Langdon, E. M., Simon-Vermot, T., Yin, A., Lee, W., Attiyeh, M., Elemento, O. and White, R. M. (2017). Melanoma genome evolution across species. BMC Genomics 18, 136.

Kirchberger, S., Sturtzel, C., Pascoal, S. and Distel, M. (2017). Quo natas, Danio?-Recent Progress in Modeling Cancer in Zebrafish. Front Oncol 7, 186.

Konantz, M., Balci, T. B., Hartwig, U. F., Dellaire, G., Andre, M. C., Berman, J. N. and Lengerke, C. (2012). Zebrafish xenografts as a tool for in vivo studies on human cancer. Hematopoietic Stem Cells Viii 1266, 124–137.

Konirova, J., Oltova, J., Corlett, A., Kopycinska, J., Kolar, M., Bartunek, P. and Zikova, M. (2017). Modulated DISP3/PTCHD2 expression influences neural stem cell fate decisions. Sci Rep 7, 41597.

Lam, S. H., Wu, Y. L., Vega, V. B., Miller, L. D., Spitsbergen, J., Tong, Y., Zhan, H., Govindarajan, K. R., Lee, S., Mathavan, S. et al. (2006). Conservation of gene expression signatures between zebrafish and human liver tumors and tumor progression. Nat Biotechnol 24, 73–5.

Love, M. I., Huber, W. and Anders, S. (2014). Moderated estimation of fold change and dispersion for RNA-seq data with DESeq2. Genome Biol 15, 550.

MacRae, C. A. and Peterson, R. T. (2015). Zebrafish as tools for drug discovery. Nat Rev Drug Discov 14, 721–31.

Manni, I., de Latouliere, L., Gurtner, A. and Piaggio, G. (2019). Transgenic Animal Models to Visualize Cancer-Related Cellular Processes by Bioluminescence Imaging. Front Pharmacol 10, 235.

McCune, J. M., Namikawa, R., Kaneshima, H., Shultz, L. D., Lieberman, M. and Weissman, I. L. (1988). The SCID-hu mouse: murine model for the analysis of human hematolymphoid differentiation and function. Science 241, 1632–9.

Miwatashi, S., Arikawa, Y., Kotani, E., Miyamoto, M., Naruo, K., Kimura, H., Tanaka, T., Asahi, S. and Ohkawa, S. (2005). Novel inhibitor of p38 MAP kinase as an anti-TNF-alpha drug: discovery of N-[4-[2-ethyl-4-(3-methylphenyl)-1,3-thiazol-5-yl]-2-pyridyl]benzamide (TAK-715) as a potent and orally active anti-rheumatoid arthritis agent. J Med Chem 48, 5966–79.

Moore, J. C., Tang, Q., Yordan, N. T., Moore, F. E., Garcia, E. G., Lobbardi, R., Ramakrishnan, A., Marvin, D. L., Anselmo, A., Sadreyev, R. I. et al. (2016). Single-cell imaging of normal and malignant cell engraftment into optically clear prkdc-null SCID zebrafish. J Exp Med 213, 2575–2589.

Mosimann, C., Kaufman, C. K., Li, P., Pugach, E. K., Tamplin, O. J. and Zon, L. I. (2011). Ubiquitous transgene expression and Cre-based recombination driven by the ubiquitin promoter in zebrafish. Development 138, 169–77.

Oltova, J., Jindrich, J., Skuta, C., Svoboda, O., Machonova, O. and Bartunek, P. (2018). Zebrabase: An Intuitive Tracking Solution for Aquatic Model Organisms. Zebrafish 15, 642–647.

Oprisoreanu, A. M., Smith, H. L., Krix, S., Chaytow, H., Carragher, N. O., Gillingwater, T. H., Becker, C. G. and Becker, T. (2021). Automated in vivo drug screen in zebrafish identifies synapse-stabilising drugs with relevance to spinal muscular atrophy. Dis Model Mech 14.

Patro, R., Duggal, G., Love, M. I., Irizarry, R. A. and Kingsford, C. (2017). Salmon provides fast and bias-aware quantification of transcript expression. Nat Methods 14, 417–419.

Patton, E. E., Mueller, K. L., Adams, D. J., Anandasabapathy, N., Aplin, A. E., Bertolotto, C., Bosenberg, M., Ceol, C. J., Chi, P., Herlyn, M. et al. (2021a). Melanoma models for the next generation of therapies. Cancer Cell.

Patton, E. E., Widlund, H. R., Kutok, J. L., Kopani, K. R., Amatruda, J. F., Murphey, R. D., Berghmans, S., Mayhall, E. A., Traver, D., Fletcher, C. D. et al. (2005). BRAF mutations are sufficient to promote nevi formation and cooperate with p53 in the genesis of melanoma. Curr Biol 15, 249–54.

Patton, E. E., Zon, L. I. and Langenau, D. M. (2021b). Zebrafish disease models in drug discovery: from preclinical modelling to clinical trials. Nat Rev Drug Discov.

Potts, K. S. and Bowman, T. V. (2017). Modeling Myeloid Malignancies Using Zebrafish. Front Oncol 7, 297.

Povoa, V., Rebelo de Almeida, C., Maia-Gil, M., Sobral, D., Domingues, M., Martinez-Lopez, M., de Almeida Fuzeta, M., Silva, C., Grosso, A. R. and Fior, R. (2021). Innate immune evasion revealed in a colorectal zebrafish xenograft model. Nat Commun 12, 1156.

Precazzini, F., Pancher, M., Gatto, P., Tushe, A., Adami, V., Anelli, V. and Mione, M. C. (2020). Automated in vivo screen in zebrafish identifies Clotrimazole as targeting a metabolic vulnerability in a melanoma model. Dev Biol 457, 215–225.

Pruitt, M. M., Marin, W., Waarts, M. R. and de Jong, J. L. O. (2017). Isolation of the Side Population in Myc-induced T-cell Acute Lymphoblastic Leukemia in Zebrafish. J Vis Exp.

Pruvot, B., Jacquel, A., Droin, N., Auberger, P., Bouscary, D., Tamburini, J., Muller, M., Fontenay, M., Chluba, J. and Solary, E. (2011). Leukemic cell xenograft in zebrafish embryo for investigating drug efficacy. Haematologica 96, 612–6.

Rheault, T. R., Donaldson, K. H., Badiang-Alberti, J. G., Davis-Ward, R. G., Andrews, C. W., Bambal, R., Jackson, J. R. and Cheung, M. (2010). Heteroaryl-linked 5-(1H-benzimidazol-1-yl)-2-thiophenecarboxamides: potent inhibitors of polo-like kinase 1 (PLK1) with improved drug-like properties. Bioorg Med Chem Lett 20, 4587–92.

Ribas, A., Lawrence, D., Atkinson, V., Agarwal, S., Miller, W. H., Carlino, M. S., Fisher, R., Long, G. V., Hodi, F. S., Tsoi, J. et al. (2019). Combined BRAF and MEK inhibition with PD-1 blockade immunotherapy in BRAF-mutant melanoma. Nature Medicine 25, 936–940.

Ridges, S., Heaton, W. L., Joshi, D., Choi, H., Eiring, A., Batchelor, L., Choudhry, P., Manos, E. J., Sofla, H., Sanati, A. et al. (2012). Zebrafish screen identifies novel compound with selective toxicity against leukemia. Blood 119, 5621–31.

Richter, S., Schulze, U., Tomancak, P. and Oates, A. C. (2017). Small molecule screen in embryonic zebrafish using modular variations to target segmentation. Nat Commun 8, 1901.

Robert, C., Grob, J. J., Stroyakovskiy, D., Karaszewska, B., Hauschild, A., Levchenko, E., Chiarion Sileni, V., Schachter, J., Garbe, C., Bondarenko, I. et al. (2019). Five-Year Outcomes with Dabrafenib plus Trametinib in Metastatic Melanoma. N Engl J Med 381, 626–636.

Rocca, S., Carra, G., Poggio, P., Morotti, A. and Brancaccio, M. (2018). Targeting few to help hundreds: JAK, MAPK and ROCK pathways as druggable targets in atypical chronic myeloid leukemia. Mol Cancer 17, 40.

Sabbah, M., Najem, A., Krayem, M., Awada, A., Journe, F. and Ghanem, G. E. (2021). RTK Inhibitors in Melanoma: From Bench to Bedside. Cancers (Basel) 13.

Schaub, F. X., Reza, M. S., Flaveny, C. A., Li, W., Musicant, A. M., Hoxha, S., Guo, M., Cleveland, J. L. and Amelio, A. L. (2015). Fluorophore-NanoLuc BRET Reporters Enable Sensitive In Vivo Optical Imaging and Flow Cytometry for Monitoring Tumorigenesis. Cancer Res 75, 5023–33.

Schindelin, J., Arganda-Carreras, I., Frise, E., Kaynig, V., Longair, M., Pietzsch, T., Preibisch, S., Rueden, C., Saalfeld, S., Schmid, B. et al. (2012). Fiji: an open-source platform for biological-image analysis. Nature Methods 9, 676–682.

Skuta, C., Popr, M., Muller, T., Jindrich, J., Kahle, M., Sedlak, D., Svozil, D. and Bartunek, P. (2017). Probes &Drugs portal: an interactive, open data resource for chemical biology. Nat Methods 14, 759–760.

Somasagara, R. R., Huang, X., Xu, C., Haider, J., Serody, J. S., Armistead, P. M. and Leung, T. (2021). Targeted therapy of human leukemia xenografts in immunodeficient zebrafish. Sci Rep 11, 5715.

Sonoshita, M., Scopton, A. P., Ung, P. M. U., Murray, M. A., Silber, L., Maldonado, A. Y., Real, A., Schlessinger, A., Cagan, R. L. and Dar, A. C. (2018). A whole-animal platform to advance a clinical kinase inhibitor into new disease space. Nat Chem Biol 14, 291–298.

Stacer, A. C., Nyati, S., Moudgil, P., Iyengar, R., Luker, K. E., Rehemtulla, A. and Luker, G. D. (2013). NanoLuc reporter for dual luciferase imaging in living animals. Mol Imaging 12, 1–13.

Takai, N., Hamanaka, R., Yoshimatsu, J. and Miyakawa, I. (2005). Polo-like kinases (Plks) and cancer. Oncogene 24, 287–91.

Tambe, M., Karjalainen, E., Vaha-Koskela, M., Bulanova, D., Gjertsen, B. T., Kontro, M., Porkka, K., Heckman, C. A. and Wennerberg, K. (2020). Pan-RAF inhibition induces apoptosis in acute myeloid leukemia cells and synergizes with BCL2 inhibition. Leukemia 34, 3186–3196.

Tang, Q., Abdelfattah, N. S., Blackburn, J. S., Moore, J. C., Martinez, S. A., Moore, F. E., Lobbardi, R., Tenente, I. M., Ignatius, M. S., Berman, J. N. et al. (2014). Optimized cell transplantation using adult rag2 mutant zebrafish. Nat Meth 11, 821–824.

Tang, Q., Moore, J. C., Ignatius, M. S., Tenente, I. M., Hayes, M. N., Garcia, E. G., Torres Yordan, N., Bourque, C., He, S., Blackburn, J. S. et al. (2016). Imaging tumour cell heterogeneity following cell transplantation into optically clear immune-deficient zebrafish. Nat Commun 7.

Tobia, C., Coltrini, D., Ronca, R., Loda, A., Guerra, J., Scalvini, E., Semeraro, F. and Rezzola, S. (2021). An Orthotopic Model of Uveal Melanoma in Zebrafish Embryo: A Novel Platform for Drug Evaluation. Biomedicines 9.

Topczewska, J. M., Postovit, L. M., Margaryan, N. V., Sam, A., Hess, A. R., Wheaton, W. W., Nickoloff, B. J., Topczewski, J. and Hendrix, M. J. (2006). Embryonic and tumorigenic pathways converge via Nodal signaling: role in melanoma aggressiveness. Nat Med 12, 925–32.

Tulotta, C., Stefanescu, C., Beletkaia, E., Bussmann, J., Tarbashevich, K., Schmidt, T. and Snaar-Jagalska, B. E. (2016). Inhibition of signaling between human CXCR4 and zebrafish ligands by the small molecule IT1t impairs the formation of triple-negative breast cancer early metastases in a zebrafish xenograft model. Dis Model Mech 9, 141–53.

Tulotta, C., Stefanescu, C., Chen, Q., Torraca, V., Meijer, A. H. and Snaar-Jagalska, B. E. (2019). CXCR4 signaling regulates metastatic onset by controlling neutrophil motility and response to malignant cells. Sci Rep 9, 2399.

Vetrie, D., Helgason, G. V. and Copland, M. (2020). The leukaemia stem cell: similarities, differences and clinical prospects in CML and AML. Nat Rev Cancer 20, 158–173.

Wertman, J., Veinotte, C. J., Dellaire, G. and Berman, J. N. (2016). The Zebrafish Xenograft Platform: Evolution of a Novel Cancer Model and Preclinical Screening Tool. Adv Exp Med Biol 916, 289–314.

Westerfield, M. (2007). THE ZEBRAFISH BOOK: A guide for the laboratory use of zebrafish (Danio rerio), 5th Edition. Eugene, University of Oregon Press.

White, R., Rose, K. and Zon, L. (2013). Zebrafish cancer: the state of the art and the path forward. Nat Rev Cancer 13, 624–36.

White, R. M., Cech, J., Ratanasirintrawoot, S., Lin, C. Y., Rahl, P. B., Burke, C. J., Langdon, E., Tomlinson, M. L., Mosher, J., Kaufman, C. et al. (2011). DHODH modulates transcriptional elongation in the neural crest and melanoma. Nature 471, 518–22.

White, R. M., Sessa, A., Burke, C., Bowman, T., LeBlanc, J., Ceol, C., Bourque, C., Dovey, M., Goessling, W., Burns, C. E. et al. (2008). Transparent adult zebrafish as a tool for in vivo transplantation analysis. Cell Stem Cell 2, 183–9.

Wickham, H., Averick, M., Bryan, J., Chang, W., McGowan, L., François, R., Grolemund, G., Hayes, A., Henry, L., Hester, J. et al. (2019). Welcome to the Tidyverse. Journal of Open Source Software 4.

Yan, C., Brunson, D. C., Tang, Q., Do, D., Iftimia, N. A., Moore, J. C., Hayes, M. N., Welker, A. M., Garcia, E. G., Dubash, T. D. et al. (2019). Visualizing Engrafted Human Cancer and Therapy Responses in Immunodeficient Zebrafish. Cell.

Yan, C., Do, D., Yang, Q., Brunson, D. C., Rawls, J. F. and Langenau, D. M. (2020). Single-cell imaging of human cancer xenografts using adult immunodeficient zebrafish. Nat Protoc 15, 3105–3128.

Yen, J. H., Lin, C. Y., Chuang, C. H., Chin, H. K., Wu, M. J. and Chen, P. Y. (2020). Nobiletin Promotes Megakaryocytic Differentiation through the MAPK/ERK-Dependent EGR1 Expression and Exerts Anti-Leukemic Effects in Human Chronic Myeloid Leukemia (CML) K562 Cells. Cells 9.

